# Validation of Structure Tensor Analysis for Orientation Estimation in Brain Tissue Microscopy

**DOI:** 10.1101/2025.01.16.633408

**Authors:** Bryson Gray, Andrew Smith, Allan MacKenzie-Graham, David W. Shattuck, Daniel Tward

## Abstract

Accurate localization of white matter pathways using diffusion MRI is critical to investigating brain connectivity, but the accuracy of current methods is not thoroughly understood. A fruitful approach to validating accuracy is to consider microscopy data that have been co-registered with MRI of post mortem samples. In this setting, structure tensor analysis is a standard approach to computing local orientations for validation. However, structure tensor analysis itself has not been well-validated and is subject to uncertainty in its angular resolution, and selectivity to specific spatial scales. In this work, we conducted a simulation study to investigate the accuracy of using structure tensors to estimate the orientations of fibers arranged in configurations with and without crossings. We examined a range of simulated conditions, with a focus on investigating the method’s behavior on images with anisotropic resolution, which is particularly common in microscopy data acquisition. We also analyzed 2D and 3D optical microscopy data. Our results show that parameter choice in structure tensor analysis has relatively little effect on accuracy for estimating single orientations, although accuracy decreases with anisotropy. On the other hand, when estimating the orientations of crossing fibers, the choice of parameters becomes critical, and poor choices result in orientation estimates that are essentially random. This work provides a set of recommendations for researchers seeking to apply structure tensor analysis effectively in the study of axonal orientations in brain imaging data and quantifies the method’s limitations, particularly in the case of anisotropic data.

**Highlights:**

- Structure tensor estimates of homogeneous orientations are robust to parameter choice
- Accuracy in crossing-fiber regions is highly sensitive to parameter choice
- Image anisotropy has a large effect on accuracy in 3D images
- Simulations provide guidelines for choosing parameters based on image characteristics

## 1. Introduction

Over the past few decades, the scope of neuroscience research has broadened from a focus on mapping the functions of brain areas with increasingly fine resolution to investigating brain networks, which are believed to be mediated by white matter (WM), the myelinated portion of the brain that makes up the projections between cortical regions. This shift is supported by accumulated evidence showing that differences in WM correlate with a wide range of neurological and psychiatric diseases and disorders, including Alzheimer’s disease, autism spectrum disorder, major depressive disorder, bipolar disorder, and schizophrenia (Fields, 2008). Two recent large-scale research initiatives to map the brain’s connectome exemplify this change in research focus: the Human Connectome Project and the BRAIN Initiative Connectivity Across Scales (BRAIN CONNECTS) program. These studies rely heavily on diffusion magnetic resonance imaging (dMRI). By measuring the orientation of water diffusion, dMRI can be used to generate estimated reconstructions of the pathways traversed by WM bundles using any of the various computational tractography methods (Smith et al., 2020). It is currently the only noninvasive method available for this purpose. Accurate reconstruction of WM pathways is also an important component of clinical research and practice. Studies have shown improved patient outcomes after brain surgery when dMRI tractography was used to locate WM bundles for surgical planning and navigation to avoid interference with critical pathways (Toescu et al., 2021). However, a scarcity of ground-truth data for validating the correspondence between diffusion models based on dMRI and true axon orientations presents a challenge for interpreting WM reconstructions derived from dMRI. In fact, a study that applied multiple methods to reconstruct simulated WM bundles observed that computational tractography studies may identify four times as many false positive connections as they do true positive connections (Maier-Hein et al., 2017).

Several studies have used structure tensor (ST) analysis—an image processing method for extracting orientation information from images—to estimate the orientation of axons in brain tissue microscopy and compare these to orientation estimates derived from dMRI. Because ST analysis is applied to images with much higher resolutions than dMRI, some authors have considered it to be the ground truth in these comparisons. Budde and Frank (2012) and Budde and Annese (2013) first demonstrated the potential for ST analysis of histologically stained sections to validate dMRI. They showed the relationship between low-resolution imaging features and true fiber distributions, including examples of intersecting fibers and partial volume effect produced by orthogonal adjacent fiber populations. They also showed a high correlation between angular anisotropy derived from ST analysis and dMRI. Mitter et al. (2015) applied ST analysis to post mortem histology of human fetal brains to validate abnormalities found *in utero* with diffusion tensor imaging. However, these studies were limited to 2D histology to validate 3D dMRI. Other groups have extended ST analysis to three dimensions for more direct comparisons. Khan et al. (2015) describe a procedure for ST analysis of z-stacks of confocal images and show that the derived orientations correspond well with the diffusion tensors in three regions of tissue with mutually orthogonal fiber directions. Schilling et al. (2016) used a similar procedure with confocal microscopy data to compare dMRI orientation distributions with true fiber distributions. They observed angular errors of 6 degrees in regions with parallel fibers and 10-11 degrees in regions with crossing fibers. ST analysis was also considered to be the ground truth for quantifying and comparing the accuracy of several different high angular resolution diffusion imaging models (Schilling et al., 2018) and to measure the fiber response function for spherical deconvolution in dMRI orientation estimation (Schilling et al., 2019). These studies inform important decisions when modeling orientation in dMRI. Thus, the sensitivity to ST analysis parameter choices and confidence estimates of the derived orientations will likely be of interest in future studies.

ST analysis was first formulated by Bigun and Granlund (1987) to detect orientations in images. The method uses partial derivative convolution filters to compute local image gradients that constitute the components of the structure tensors. A smoothing kernel is applied to the gradients to average over a local neighborhood. The eigenvectors corresponding to the smallest eigenvalues of the tensors point in the direction of the minimum gradient, i.e., parallel to the fiber directions. This method requires two parameters to be selected: the standard deviation for an isotropic first-derivative Gaussian filter (*σ*_*g*_) or pre-smoothing kernel, which controls the size of the structure to which the kernel is sensitive, and the standard deviation of the window function (*σ*_*w*_), which controls the size of the neighborhood that is blurred. These parameters were described as being analogous to the diffusion imaging time and the dMRI resolution, respectively (Khan et al., 2015). ST analysis is popular in part because of its computational efficiency relative to other methods such as higher-order derivatives, Fourier methods, directional wavelets and others as discussed in Püspöki et al. (2016). However, it has a weakness in that using a fixed pair of filter widths for the entire image limits its sensitivity to one spatial scale for an entire image (Püspöki et al., 2016). The filter output is highly sensitive to the filter parameters and the size of the structure being analyzed. If the kernel size is too small, it becomes oversensitive to noise and measures isotropic orientations inside fiber boundaries instead of characterizing the entire fiber by a single orientation. Conversely, if the kernel size is too large, it may blur orientation information across multiple fibers. For this reason, and because they observed a bias toward detecting two orientations in isotropic regions, Ning et al. (2021) recommended that care be taken when using structure tensors to identify fiber crossings. A separate study compared ST orientations with optic axis orientation measurements from optical coherence scanner images, finding agreement within 10 degrees in well-delineated fibers (Wang et al., 2015). A study by Schilling et al. (2016) included an analysis of the sensitivity of structure tensors to Gaussian filter standard deviations and in-plane resolution. They derived STs from 3D confocal microscopy with high resolutions (between 0.08 µm and 0.42 µm) and compared ST derived orientations to 100 manually traced fibers in three single-fiber regions. They reported accuracy within 5 degrees, which was relatively robust to the standard deviation of the spatial averaging filter but highly sensitive to the standard deviation of the derivative filter. While this accuracy appears relatively good, other authors Leuze et al. (2021) have found angular errors as high as 45 degrees in some regions. Schilling et al. (2016) observed that derivative filters with standard deviations close to the size of myelinated fibers (0.5 µm and 1.5 µm) produced the highest accuracy. However, the studies by Wang et al. (2015) and Schilling et al. (2016) were both performed at relatively high resolution and were limited to well-delineated fiber regions. It remains unclear how the standard deviation parameters affect the accuracy of the orientation estimates derived using ST analysis in regions with crossing fibers, or what the limits of accuracy are as the resolution decreases and anisotropy increases. This is an important question to address because the point spread function of a confocal microscope is known to have a depth nearly three times its width (Pawley, 2006), and 2D microscopy imaging for whole brain mapping studies is often restricted to slices spaced more than 50 µm apart (Amato et al., 2016; Oh et al., 2014; Ragan et al., 2012).

We believe that a more thorough understanding of the effects of parameter choice and of accuracy limitations, both of which are currently unknown, will be useful for researchers with data of differing dimensions and qualities and ascribe a confidence level to their ST analysis. We addressed these unknowns by producing digital phantoms that simulate brain histology images with different line patterns, including crossing fibers, and different degrees of image anisotropy. We then computed STs from these phantoms and quantified the accuracy of varying filter parameters. Additionally, we performed structure tensor analysis in 2D and 3D regions that we selected manually from mouse brain microscopy images that included intersecting axons. We visualized the resulting orientation distributions to provide a qualitative evaluation of the effects of parameter choice in real images.

## 2. Methods

We generated a series of 2D and 3D image data designed to simulate optical microscopy acquisitions of parallel or crossing fibers. Specifically, these digital phantoms consisted of grids of parallel lines, repeated at a specified period. These lines were rendered at a specific angle relative to one image axis for parallel fibers, or at two specific angles for crossing fibers. The image resolution along the first axis was varied to create anisotropic images of a fixed total size. We evaluated two widely-employed techniques for anisotropy correction: upsampling by linear interpolation to isotropic pixel sizes and upsampling with isotropic blur correction, where additional blur is introduced along the high-resolution directions to create equal blur along each dimension. The period (i.e., spacing between lines) and angles were also varied to simulate the variety of fiber arrangements found in tissue micrographs.

We analyzed our phantom images by computing structure tensors at every image voxel and deriving local image orientations from the eigenvectors of the structure tensors. Finally, we clustered the resulting orientations to extract the dominant estimated orientations from the image and compared them with the true orientations to quantify accuracy. We repeated this procedure for a range of *σ*_*g*_ and *σ*_*w*_ values. This method of aggregating orientations computed at high resolution to obtain the dominant orientations per image region simulates the procedure used in the studies described above to compare microscopy imaging data with lower-resolution dMRI (Budde and Frank, 2012; Budde and Annese, 2013; Mitter et al., 2015; Khan et al., 2015; Schilling et al., 2016, 2018, 2019). Accuracy results were averaged across all angles and periods in the results we show because, in general, fiber orientations will be unknown before a sample is placed in an imaging system. In the following sections, we describe in detail each component of our validation pipeline, which is summarized in Fig. 1.

**Figure 1:**
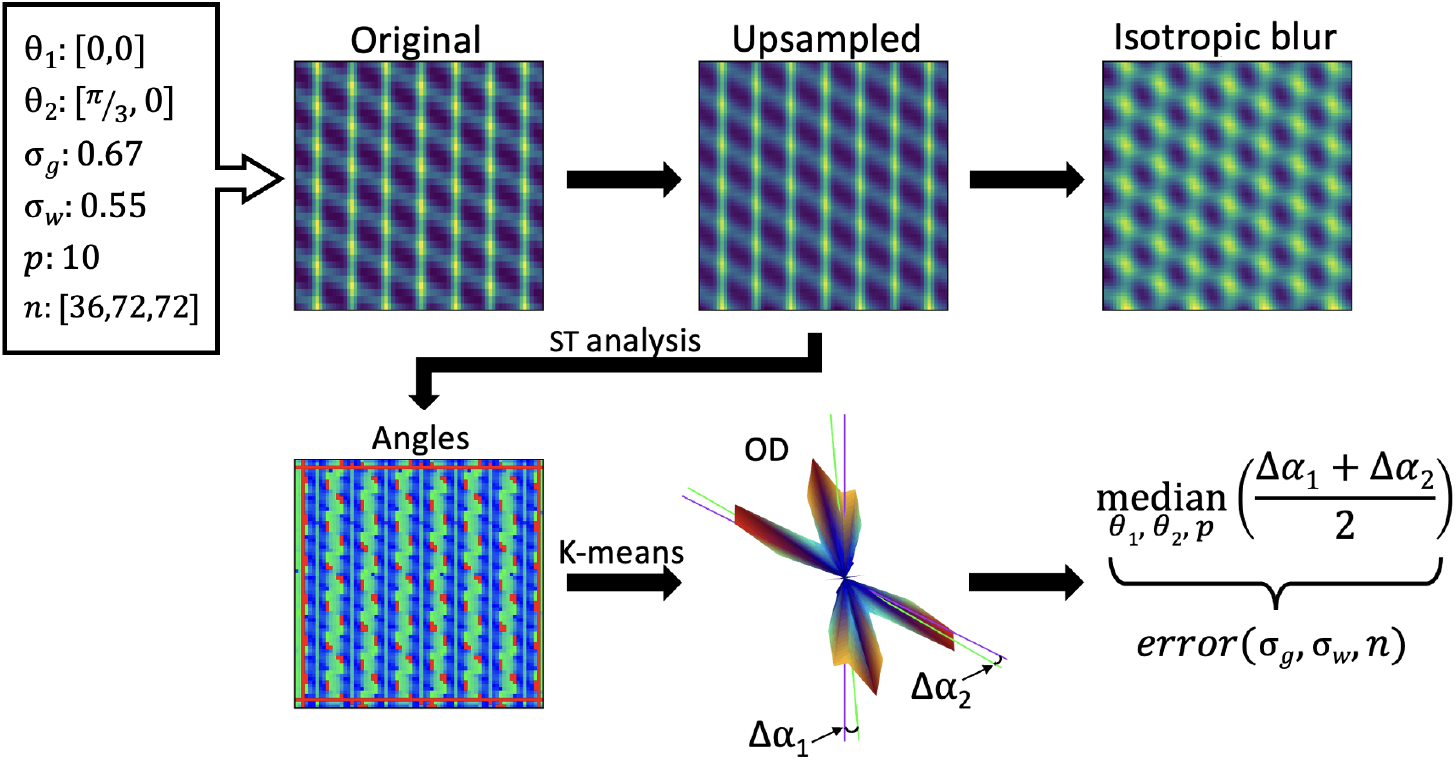
Structure tensor validation pipeline. Example pipeline using a phantom with two angles and an anisotropy ratio of 2.0. Each line phantom is specified by its angle(s) (*θ*), period (*p*), and number of pixels in each dimension (*n*). In this example, ST analysis is performed on the phantom after upsampling but not isotropic blur correction. ST analysis is parameterized by gradient and window standard deviations, *σ*_*g*_ and *σ*_*w*_. The boundaries of the images are cropped to remove boundary artifacts. The edges after cropping are shown in red. The orientation distribution (OD) represents the histogram of orientations as amplitude on a sphere. Computed means are represented by green lines through the OD with true orientations in purple. In the final step, we compute the median error over period and angles to form error as a function of standard deviations and anisotropy.

### 2.1. Phantom Construction

Each phantom fiber was represented as an ideal line convolved with a Gaussian point spread function, *k*. This Gaussian has a diagonal covariance with one anisotropic direction. In 2D, we denote the standard deviation in the high-resolution direction as *σ*_1_, and that in the low-resolution direction as *σ*_2_. In 3D, we denote both standard deviations in the two high-resolution directions as *σ*_1_ and the standard deviation in the low-resolution direction as *σ*_2_. In our analysis, we chose these parameters equal to the pixel size, which was different in both directions. The idea here is that, generally, pixel sizes for sampling are chosen to be similar to the resolution of the imaging system along each axis.

We model an axon as an infinitely thin line *ℓ*, oriented in the direction *a* ∈ ℝ^*d*^ (*d* = 1 or 2) with |*a*| = 1, with offset *b* ∈ ℝ^*d*^. We define

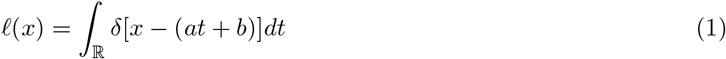

where *δ* is the Dirac delta distribution. We define our image *I* by convolving with *k* as

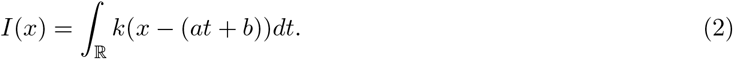

In 2D, we can derive an expression for the standard deviation of the Gaussian normal to a straight line oriented along an angle *θ*: 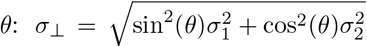. In this case, we use an analytical formula to produce lines. In 3D, no such simplification exists. Instead, we define a Gaussian function analytically and evaluate it at a rotated set of points. In all cases, we use modular arithmetic on the pixel locations to create a periodic pattern efficiently. We constructed all phantoms by sampling these analytical functions on a regular voxel grid, with 96 pixels in the high-resolution direction in 2D or 72 pixels in the high-resolution directions in 3D. We varied the number of pixels in the low-resolution direction based on the anisotropy factor to give a square or cube. After line drawing, we resampled to an isotropic grid using one of two approaches. In the first approach, we upsampled the phantoms along the low-resolution dimension using linear interpolation to create isotropic voxel dimensions while maintaining an anisotropic blur. For the second approach, we created a second set of phantoms in the same way, including upsampling to isotropic voxel dimensions, but with the addition of Gaussian blur along the high-resolution dimension(s) with variance equal to the difference of the squared pixel sizes in the lowand high-resolution directions (resulting in isotropic pixel size and isotropic blur). This step has been applied in at least one previous study for anisotropy correction (Khan et al., 2015).

Line directions were specified using a single angle for the 2D case or two angles for the 3D case. For a single angle in two dimensions, we chose to use 100 angles spaced evenly over the interval [0, *π/*2). We do not include angles outside this range because the symmetry of the phantom makes them redundant. In three dimensions, we chose 100 angles distributed on a hemisphere using an electrostatic dispersion algorithm (Jones et al., 1999), implemented in Python as the disperse charges method in Dipy (Garyfallidis et al., 2014), using 2000 iterations and a step size constant of 0.1. For phantoms with two fiber directions, in both the 2D and 3D cases, given a set of 100 evenly spaced angles, 𝒮, we defined the set of all possible pairs of angles, 𝒫 = {(*a, b*) | *a, b* ∈ 𝒮 and *a* = *b*}, and constructed a set of phantoms using a random sample without replacement of 100 angle pairs from 𝒫. This method represents the full range of crossing fibers that may be encountered in a real brain. Several examples of 2D crossing-line phantoms with varying anisotropy and line angles are shown in Figure 2. One important feature of our phantoms is that lines parallel to the high-resolution direction appear dimmer because their signal power is spread over a longer distance.

**Figure 2:**
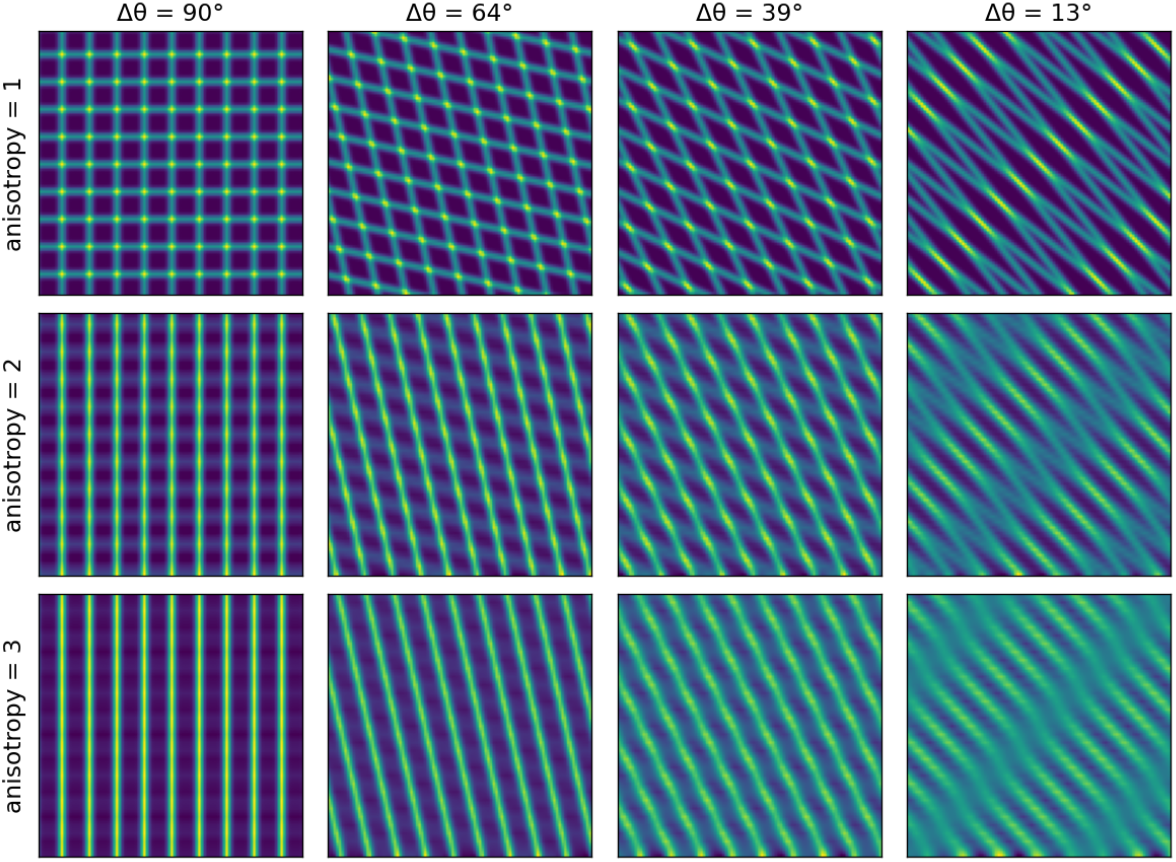
Example Phantoms. Examples of 2D crossing-line phantoms showing the effect of varying anisotropy and angle values.

We recognized that the phantoms described above represent fluorescent microscopy images well, but the summation of crossing lines creates a pattern that could affect the measured orientation distribution and that this artifact would be less prominent in bright-field microscopy. We modeled bright-field images by modifying the original phantoms to represent exponential attenuation of photons according to the Beer-Lambert law as follows:

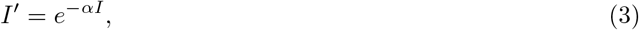

where *I* is the original phantom (a blurred bright line on a dark background), *I*^*′*^ is the bright-field type phantom, and *α* = 10 (chosen empirically to produce realistic looking images). We performed the same analysis as described above on these modified phantoms but found only very minor differences in error compared to unmodified phantoms, Thus, to avoid redundancy, we only show results from the first analysis in the subsequent sections.

### 2.2. Estimating Orientation

Structure tensor analysis begins by choosing standard deviations for the first derivative Gaussian filter and the window function. Gradients, *I*_*i*_, are then computed along each image axis *i* by applying a Gaussian filter to the image with degree one along *i* and zero along the other axes, using the same standard deviation for each axis. Because of intensity inhomogeneity and a large variety of contrast mechanisms in microscopy, we normalize the gradient vector rather than weighing brighter regions more than darker regions. A 2D or 3D structure tensor is then constructed at each voxel

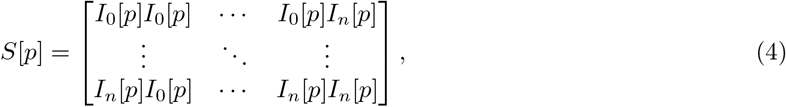

where *n* is the index of the image dimension. A Gaussian window function, *w*, is applied to the resulting matrix for local gradient averaging,

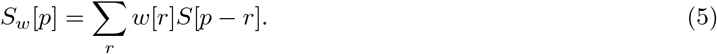

The local orientation is given by the principal eigenvector of the structure tensor. We instead acquire the direction corresponding to the smallest gradient by first subtracting the structure tensors from identity, *S*^*′*^= *Id* − *S*_*w*_. In certain cases where *S*_*w*_ was small, such as when we investigated without normalizing the gradient, we found that the eigenvalue solver we used, NumPy’s eigh function, was inaccurate. We thus opted to simply compute the eigenvalues of − *S*_*w*_. With our normalization convention, this can be thought of as a diffusion tensor under a model where at a point particles may diffuse in any direction normal to the gradient with equal probability, and parallel to the gradient with probability zero. The full-rank macroscopic diffusion tensor is found by averaging gradients near the point with weights given by *w*.

We crop the boundaries of the array of principal directions to remove boundary artifacts that are produced by interpolation and convolution. The upsampling interpolation introduces error onto one end of the low-resolution dimension of width equal to the upsampling factor minus one. We also cropped all edges by two-thirds of the radius of the largest convolution kernel. The two-thirds factor was chosen by visual inspection, because the error introduced by convolution varies by location and becomes negligible some distance away from the borders.

### 2.3. Clustering and Error Calculation

The resulting array of directions was processed using a k-means clustering procedure appropriate for antipodally symmetric directions (in this context, a direction is considered equal to its opposite) on a circle or sphere where the number of means was *k* = 1 for the single orientation experiments and *k* = 2 for the crossing-fiber experiments. The means were compared to the input angles used to generate the phantoms, and the reported error was the resultant angular difference in degrees, averaged over both angles for two-angle phantoms.

K-means clustering uses different modified distance and mean calculations for for a circle and a sphere. For a circle, the input values are angles and the distance metric is modified for periodic boundary conditions:

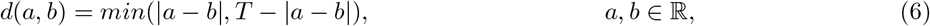

where *T* is the period; in our case, *T* = 180°. We use this distance metric to compute the Fréchet mean of the angles. That is, we find the angle, *m*, that minimizes the sum of squared distances between it and all angles in the sample:

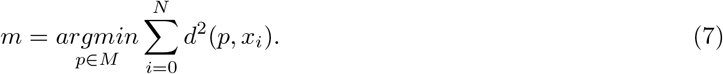

In our case, *M* is the set of whole number angles on a half-circle, {0^°^, …, 179^°^}, so that *m* is the Fréchet mean approximated to the nearest degree.

For a sphere, the input orientations are unit vectors in Cartesian coordinates. Distances are defined as

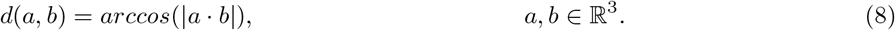

This has the range 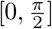. The absolute value is used to account for antipodal symmetry, that is, *d*(*x*, − *x*) = 0. The mean of the 3D direction vectors is their arithmetic mean normalized such that it lies on the unit sphere (Banerjee et al., 2005).

One modification must be made to the k-means clustering algorithm in the spherical case: if the cosine between a vector, *x*, and its assigned centroid is negative (i.e., their distance is greater than 90 degrees), the vector is reassigned to − *x*. This is necessary because the mean of two nearly antiparallel directions before normalizing is near zero even though they represent similar orientations because of symmetry. Flipping all vectors in the same cluster to the same hemisphere avoids instability from this effect. For both the 2D and 3D cases, the initial centroids are set to a random choice of *k* data points without replacement, and the stopping criterion is reached when the new centroids are equal to the old centroids.

For 2D phantoms with a single angle, we computed a single mean using the periodic mean calculation given by Eq. 7. For the two angle case, two means were computed using the periodic k-means algorithm with *k* = 2. For 3D phantoms, both one- and two-angle means were estimated using k-means clustering with *k* = 1 and *k* = 2, respectively. The distances between the computed means and the true angles were calculated using Eq. 6 for 2D or Eq. 8 for 3D. In the two-angle case the error was the average of two distances: the distance between the nearest mean and true angle pair and the distance between the remaining mean and true angle.

The resulting errors were tabulated using a Pandas data frame in Python. The error heat maps shown in Figures 4-7 were created by computing the median of the errors over period and angles and grouping by the remaining variables: *σ*_*g*_, *σ*_*w*_, and anisotropy ratio.

We validated the k-means and error calculations by computing errors using the method described above on samples from known orientation distributions. We drew samples from von Mises and von Mises-Fisher distributions using Scipy’s vonmises and vonmises fisher methods for 2D and 3D, respectively, and compared the estimated means from our methods with the true sample means over a range of variances and over varying crossing angles for mixtures of two distributions (Virtanen et al., 2020). Figure 3 shows a visual representation of a mixture of 3D von Mises-Fisher distributions and the resulting differences between k-means centers and true means.

**Figure 3:**
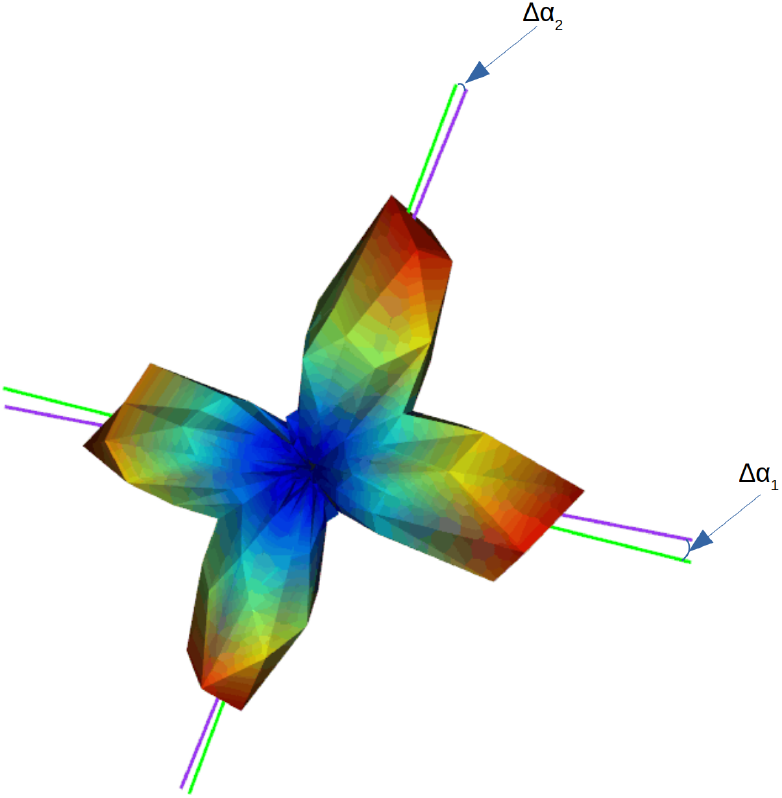
K-Means Error Calculation. The estimated means (green lines) compared to true means (purple lines) shown with a sample orientation distribution represented as a 3D histogram. The error is the mean of Δ*α*_1_ and Δ*α*_2_.

### 2.4. Example Mouse Brain Microscopy

To illustrate the applicability of our model to real data analysis, we considered image regions from mouse brain microscopy datasets and qualitatively compared the performance of structure tensor analysis with the predictions of our phantom study. We used the ImageJ software package to view 2D and 3D brain microscopy images and select two regions that exhibited two predominant intersecting axon orientations (Schindelin et al., 2012). The 2D image was acquired from a mouse brain that was fixed using cardiac perfusion (4% paraformaldehyde in 0.01M PBS) and sliced into 50 µm thick serial sections using a vibratome microtome. The slices were stained for myelin and imaged with an Olympus VS200 microscope under bright-field mode with 10X magnification at 1.1 µm *×* 1.1 µm resolution. The 3D volume was taken from an optically cleared (SHIELD, LifeCanvas Technologies, Cambridge, MA) mouse spinal cord with fluorescently labeled axons (B6.Cg-Tg(Thy1-YFP)HJrs/J, The Jackson Laboratory, Bar Harbor, ME). The spinal cord was imaged on a TCS SP8 MP multiphoton microscope (Leica Microsystems Inc., Deerfield, IL) with a Leica HC FLUOTAR L 25×/1.0 IMM objective at 0.87 µm *×* 0.87 µm *×* 2.4 µm resolution. We selected a 100 *×* 100 pixel region from the 2D image and an 812 *×* 728 *×* 29 voxel region from the 3D volume for our analysis. As with the phantoms, we applied anisotropy correction to the volume region by linear interpolation along the low-resolution dimension to isotropic pixel size, yielding a final shape of 812 *×* 728 *×* 78 after resampling and cropping the interpolation artifact. The dominant axon orientations were manually estimated for each image.

We then generated one 2D phantom and one 3D phantom, each corresponding to an example microscopy region. The axons in the example images were approximately 2 pixels wide for the 2D image and 4 pixels wide for the 3D volume. However, the phantoms in our simulation study (described in Section 2.1) simulated axons with the assumption that they were approximately one pixel wide. We created phantoms suitable for the 2D and 3D microscopy data by setting the phantom sizes and line spacings to those of the microscopy images divided by the approximate axon widths, in pixels, in those images. For each phantom, we selected angles that matched the manually estimated axon orientations. We performed ST analysis on the phantoms using the same range of *σ*_*g*_ and *σ*_*w*_ as in the phantom experiments, and we used these results to select three parameter settings to use on the example microscopy. These included one setting that predicts high accuracy and two that predict low accuracy (the selected *σ*_*g*_ and *σ*_*w*_ had to be scaled up by the axon widths to remain consistent). Microscopy image orientations, mean orientations, and errors relative to the manually annotated orientations were computed as described in Sections 2.2 and 2.3 using the selected parameters.

### 2.5. Software Availability

All software methods and example microscopy ROIs used in this paper are available under an open-source license and may be downloaded at https://github.com/BrysonGray/structure_tensor_validation.git.

## 3. Results

In the following section, we report the effects that filter parameters and image anisotropy had on the accuracy of structure tensor orientation estimates in our simulated phantoms. These results could be used to determine optimal filter widths for a given analysis task and as a guide for which suboptimal choices should be avoided. We show errors as a function of the gradient and window standard deviations and anisotropy ratio. We also report the best Gaussian filter standard deviations and the best and worst angular errors at each anisotropy value. Note that we chose to report errors based on upsampled but not isotropically blurred phantoms because the latter step reduced accuracy in all experiments. Experiments are split into single-orientation phantoms and two-orientation phantoms. Our results are the median over all line angles and periods.

### 3.1. 2D Experiments

The results of the 2D experiments can be seen in Figures 4 and 5. Note that in the single angle case, accuracy is robust to parameter choice even up to relatively high anisotropy.

**Figure 4:**
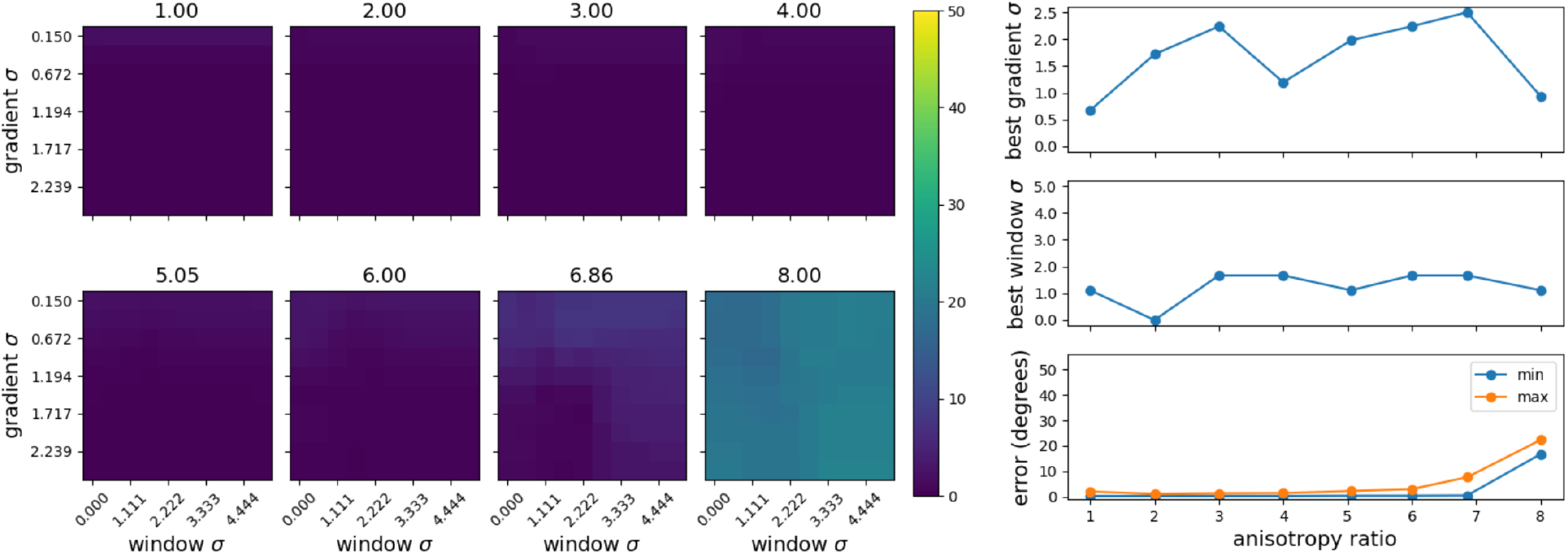
2D Single Angle. Left: Median errors in degrees with respect to *σ*_*g*_ and *σ*_*w*_ for 2D single-angle phantoms represented in a heat map with the anisotropy ratio indicated above each plot. Top right: The *σ*_*g*_ and *σ*_*w*_ values corresponding to the minimum error from each heat map. Bottom right: The minimum and maximum errors corresponding to each heat map.

**Figure 5:**
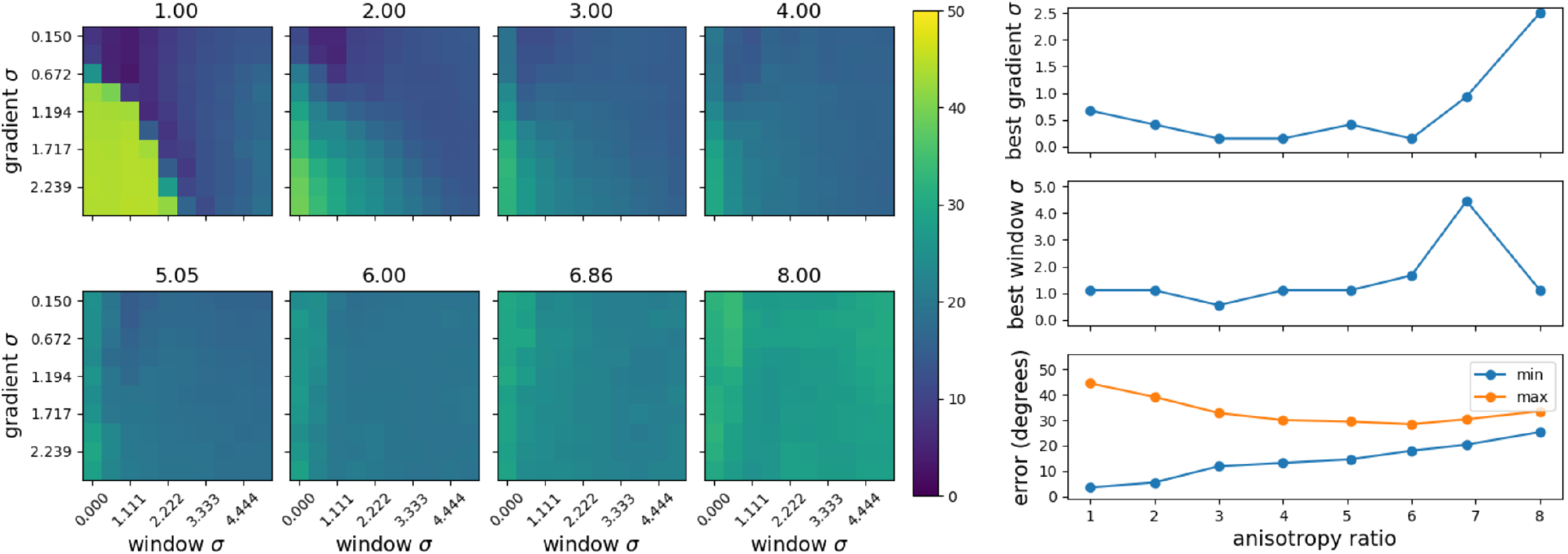
2D Two Angles. Left: Median errors in degrees with respect to *σ*_*g*_ and *σ*_*w*_ for 2D two-angle phantoms represented in a heat map with the anisotropy ratio indicated above each plot. Top right: The *σ*_*g*_ and *σ*_*w*_ values corresponding to the minimum error from each heat map. Bottom right: The minimum and maximum errors corresponding to each heat map.

Figure 5 shows a more dramatic difference in errors between good and bad parameter choices for resolving crossing fibers compared to homogeneous orientations. These results suggest that for small anisotropy ratios, it is important to choose a *σ*_*g*_ that is small relative to the fiber width and a *σ*_*w*_ that is approximately equal to the fiber width. For anisotropy ratios greater than two, accuracy becomes less sensitive to parameter choice, but decreases significantly. This decrease in sensitivity resulted in outliers in the predicted best parameters at high anisotropy, as seen in the figure.

### 3.2. 3D Experiments

Results from the 3D experiments can be seen in Figures 6 and 7. These results show trends that are similar to the 2D experiments, but with some notable differences. First, for all 3D structure tensor analysis, it is important to use *σ*_*w*_ greater than zero. This is because structure tensors are rank one prior to spatial averaging, meaning that the two smaller eigenvalues are degenerate and their eigenvectors are ambiguous. Only after spatial averaging by the window function does the tensor become full rank, and the two smaller eigenvectors become unambiguous. Note that we ignore *σ*_*w*_ equal to zero when reporting the maximum error.

**Figure 6:**
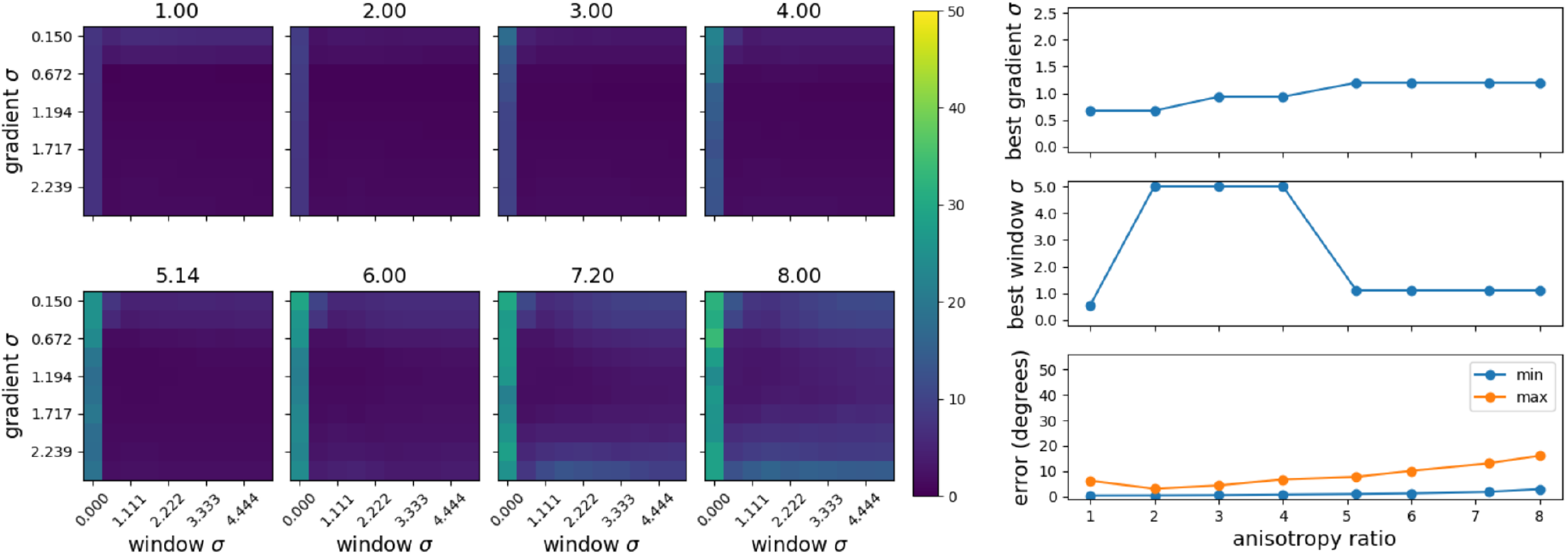
3D Single Angle. Left: Median errors in degrees with respect to *σ*_*g*_ and *σ*_*w*_ for 3D single-angle phantoms represented in a heat map with the anisotropy ratio indicated above each plot. Top right: The *σ*_*g*_ and *σ*_*w*_ values corresponding to the minimum error from each heat map. Bottom right: The minimum and maximum errors corresponding to each heat map.

**Figure 7:**
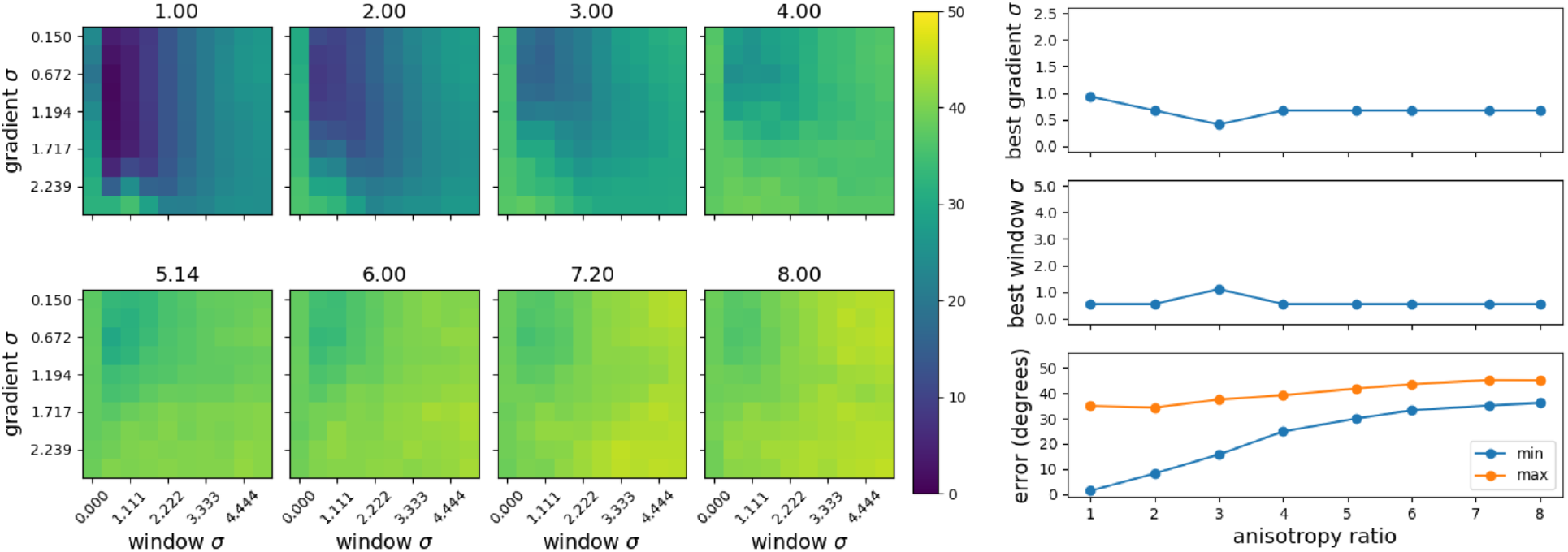
3D Two Angles. Left: Median errors in degrees with respect to *σ*_*g*_ and *σ*_*w*_ for 3D two-angle phantoms represented in a heat map with the anisotropy ratio indicated above each plot. Top right: The *σ*_*g*_ and *σ*_*w*_ values corresponding to the minimum error from each heat map. Bottom right: The minimum and maximum errors corresponding to each heat map.

Similar to the 2D case, errors are close to zero even up to large anisotropy for single angles in 3D. For any anisotropy, *σ*_*g*_ is best chosen between one and two, while *σ*_*w*_ appears to have little effect in this case.

Experiments with 3D crossing-line phantoms revealed a large difference in errors between good and bad parameters, similar to the results of the 2D crossing-line phantom experiments. Interestingly, the best parameters differ from those in the 2D case. Here, a small *σ*_*w*_ and *σ*_*g*_ near one is optimal across anisotropy settings.

### 3.3. Example Mouse Brain Microscopy Results

Figures 8 and 9 show the results of ST analysis on the example mouse brain microscopy regions alongside the manually annotated orientations and the heat map of accuracy over parameters predicted by the corresponding simulated phantom. In both the 2D and 3D cases, a qualitative comparison of the similarity between manually annotated orientations and ST derived orientations over each of three *σ* parameter settings agrees with the trends predicted by the phantom ST analysis results shown in the top-left panels. Additionally, these figures qualitatively represent the way that the ST analysis becomes oversensitive to one predominant image orientation and fails to capture intersecting orientations when either *σ*_*w*_ and *σ*_*g*_ are too large.

**Figure 8:**
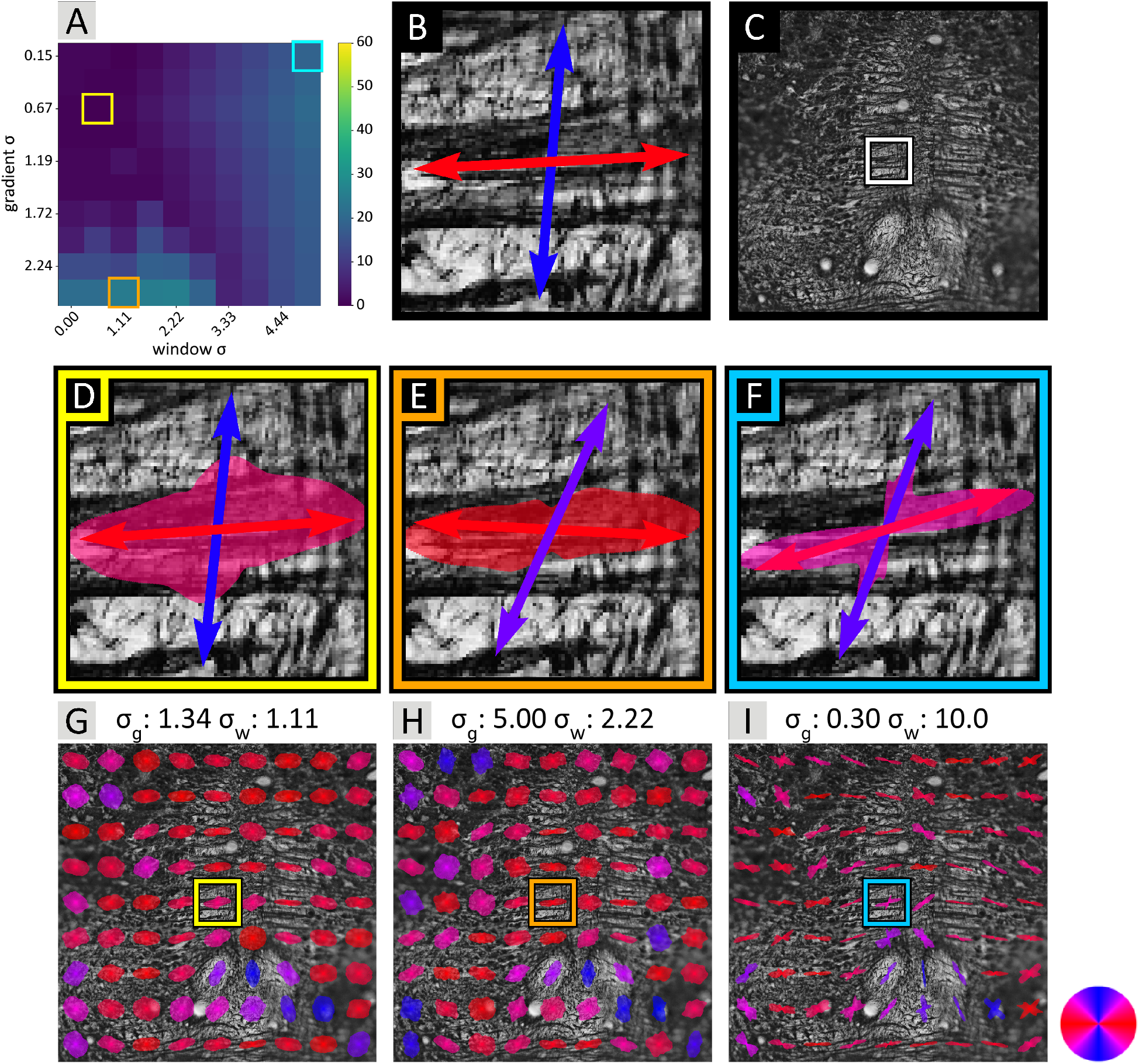
2D Example Microscopy. A bright-field image of myelin stained mouse brain is shown with dark neuron fibers visible on a light background (C). A subregion, shown as a white bounding box, was chosen for analysis. The subregion with manually identified primary fiber orientations are shown as arrows with angles mapped to colors (B). The accuracy of ST derived orientations on a matching simulated phantom is shown in the top-left (A). Three *σ* settings, indicated by colored bounding boxes, were selected from the heat map to compute orientations in the microscopy region. Note that the selected *σ*_*g*_ and *σ*_*w*_ are scaled up by two to account for the higher resolution in the example microscopy as compared to the phantoms. Derived orientations are shown as polar histograms and the computed means as arrows overlaid on the image for each *σ* setting (D-F). The same regions are shown centered in their larger context in (G-I).

**Figure 9:**
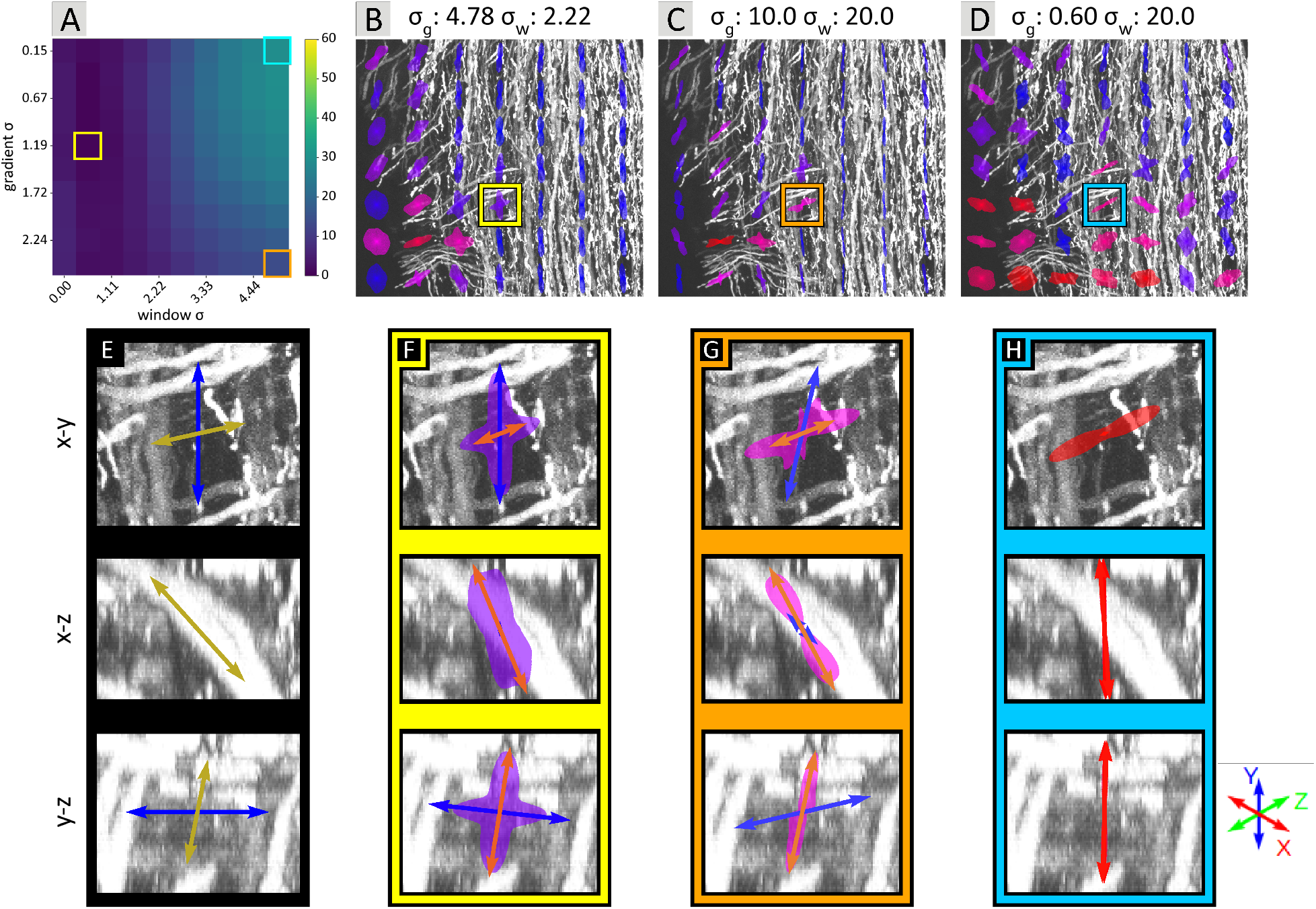
3D Example Microscopy. A small fluorescently labeled 3D volume from mouse spinal cord is shown with neurons visible as bright objects on a dark background. The volume is represented as maximum intensity projections from three orthogonal views (rows 2-4). Primary orientations are represented by arrows with angles mapped to colors and length proportional to the in-plane component of the orientation. Manually identified primary orientations are shown in (E). The accuracy of ST derived orientations on a matching simulated phantom is shown in the top-left (A). Three *σ* settings, indicated by colored bounding boxes, were selected from the heat map to compute orientations in the microscopy region. Note that the selected *σ*_*g*_ and *σ*_*w*_ are scaled up by 4 to account for the higher resolution in the example microscopy as compared to the phantoms. Derived orientations are shown as polar histograms marginalized over out-of-plane components, and the computed means as arrows overlaid on the image for each *σ* setting (F-H). The same regions are shown centered in their larger context in the x-y view in (B-D).

## 4. Discussion

In this work, we used computational phantoms to study the accuracy of ST analysis for measuring crossing and single fiber orientations in well-characterized settings. We produced a series of 2D and 3D phantoms at different anisotropy ratios, studied accuracy with different ST filter parameters, and compared them to real microscopy data. The results lead to two general conclusions. First, structure tensor analysis is potentially very accurate. For both 2D and 3D cases in regions with homogeneous orientations, structure tensors can capture angles with accuracy within 0.01 degrees in isotropic images and maintains accuracy close to one degree up to large anisotropy ratios. In regions with crossing fibers, the best accuracy for isotropic images was approximately one degree in 3D and approximately three degrees in 2D. However, it is important to be aware that this accuracy was not sustained and deteriorated as the anisotropy ratio is increased.

Second, the accuracy of structure tensor analysis in regions with crossing fibers is very sensitive to the standard deviations of the Gaussian filters used. For example, in 2D data with crossing fibers, an increase in the *σ*_*g*_ of 0.5 pixels from the optimal value reduced the accuracy from four degrees to forty degrees.

Based on these results we can make a few recommendations for researchers planning to employ structure tensor analysis. But first, we must be careful with units when discussing the values of standard deviations. Although the standard deviation is defined in units of pixels, it is more important to consider the size of the structure being analyzed when choosing these parameter values. Because the structure tensor characterizes the orientation of boundaries, using a standard deviation that is too small relative to the structure will result in isotropic orientation between the structure’s boundaries. Assigning a standard deviation that is too large causes the filter to encompass multiple structures and blur their orientation information. In our experiments, the simulated fibers were modeled as being infinitely thin lines whose measured signal is a Gaussian with width equal to a pixel, where the Gaussian is anisotropic in the case of anisotropic pixel sizes. Therefore, we will make recommendations in units of pixels (*px*), keeping in mind that we assume the structures of interest appear approximately one pixel wide in the images.

First, assuming the region of interest includes a significant amount of overlapping fibers along with regions with single orientations, our experiments suggest that high accuracy can be achieved in both regions by selecting 0.1 *px* ≤ *σ*_*g*_ ≤ 0.5 *px* and 1.0 *px* ≤ *σ*_*w*_ ≥ 2.0 *px* for 2D images and 0.5 *px* ≤ *σ*_*g*_ ≤ 1.0 *px* and 0.5 *px* ≤ *σ*_*w*_ ≤ 1.0 *px* for 3D images. We do not recommend using structure tensor analysis on images with any dimension having resolution lower than the structure being analyzed because even the best accuracy at anisotropy ratios of two or greater is below what could reasonably be considered ground truth. In addition, we do not recommend introducing blur in the x-y plane to create isotropic blur, as was implemented in (Khan et al., 2015). This only decreased the accuracy in our experiments, as shown in Figure 10. We also found that normalizing the gradients before constructing tensors had a negligible effect on accuracy. Though we did not investigate the effect of noise on accuracy, we suspect that normalizing gradients may have the undesirable effect of amplifying noise in homogeneous regions.

**Figure 10:**
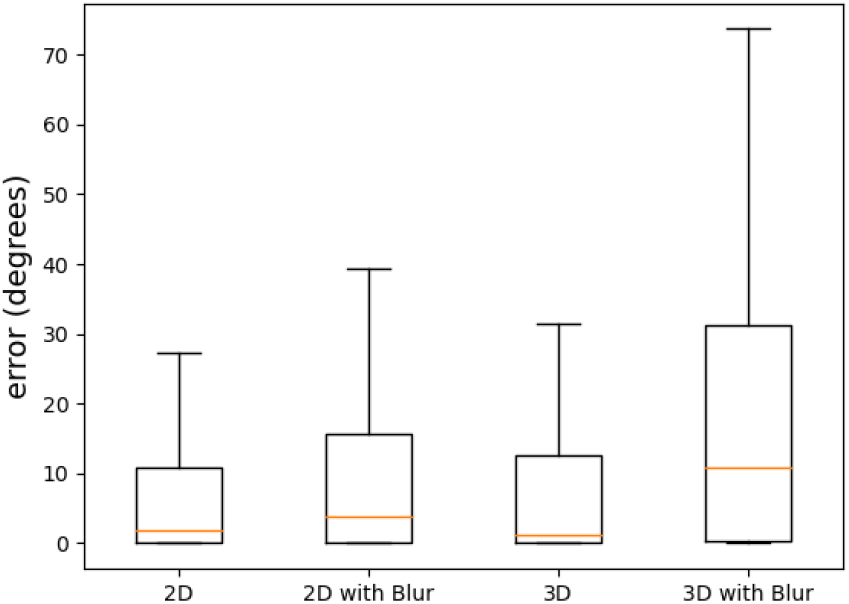
Effect of Isotropic Blur Correction. Comparison of error distributions between phantoms with isotropic blur correction and those without.

While our phantom experiments have the benefit of having known true orientations and line spacing, they come with some inherent limitations. Phantoms are idealizations of real images, which have curving lines and lines of varying thickness. With this in mind, we made an effort to identify the limits of this method and make it generalize as much as possible. We modeled both fluorescent images and bright-field images. It is notable that we observed a negligible difference between summed lines (fluorescent-type) phantoms and exponentially attenuated (bright-field-type) phantoms. This suggests that the artifact caused by summing intersecting lines does not significantly affect the errors. In our example microscopy data, both the curving of axons and the variation in thickness are quite apparent. Nevertheless, our model is able to predict example parameter choices that may lead to poor performance, and example parameter choices that lead to good performance. That being said, the phantom study results we reported were averaged across all simulated angles, whereas the microscopy datasets only have a few primary orientations. While exploring performance as a function of angle will be the subject of future work, its applicability may be limited because angular distributions are unknown until images have been acquired. For completeness, the supplementary data we share in our GitHub repository includes accuracy results as a function of angle.

The method used to aggregate structure tensors and characterize orientation in a local region is also relevant. K-means clustering is known to be sensitive to initial centers, so it is common when employing kmeans clustering to try several different initial centers and use the estimate that achieves the smallest sum of squared distance of points from their cluster centers. When testing our k-means on known distributions, we found small differences from the known means. Increasing the number of initializations did not significantly change the difference between estimated and true means so we chose to use one initialization to reduce computation time.

Finally, while the lines we drew were sufficiently anti-aliased, the details disappeared and the pattern blurred away as the anisotropy ratio approached the period of our line pattern. The smallest period we used was 7.0 and the greatest anisotropy ratio we used was 8.0, thus there was significant error due to this artifact. This can be seen in Figure 4. This demonstrates how substantially fiber orientation accuracy is compromised even with moderate anisotropy levels. The anisotropy and structure size are important considerations for researchers intending to apply ST analysis to their datasets, and good accuracy could likely be achieved by following our recommendations above.

Overall, the studies presented in this work can give confidence to investigators who consider applying ST analysis by clarifying the range of data and parameters appropriate for its use and by demonstrating the high degree of accuracy that can be achieved under the right conditions. This work will help give researchers certainty in the accuracy of their diffusion MRI images and downstream processing. We expect this will have the potential to impact development of tractography algorithms and analysis of brain connectivity. This may lead to improvements in our understanding of basic neuroscience (cell types are often understood based on their connectivity) (Hawrylycz et al., 2023) and medical applications of connectomics (Toescu et al., 2021).

## 5. Acknowledgements

We gratefully acknowledge the contribution of our collaborators at UCLA, Hongwei Dong and Hanpeng Xu, who provided 2D myelin-stained microscopy images.

## 6. Author Contributions

**Bryson Gray**: Conceptualization, Methodology, Software, Validation, Formal analysis, Investigation, Data curation, Writing Original draft, Writing Review & editing, Visualization. **Andrew Smith**: Resources, Data curation, Writing - Review & editing. **Allan MacKenzie-Graham**: Resources, Data curation, Writing - Review & editing, Supervision, Project administration, Funding acquisition. **David Shattuck**: Conceptualization, Methodology, Resources, Writing - Original draft, Writing - Review & editing, Funding acquisition. **Daniel Tward**: Conceptualization, Methodology, Software, Validation, Formal analysis, Resources, Data curation, Writing - Original draft, Writing - Review & editing, Visualization, Supervision, Project administration, Funding acquisition.

## 7. Funding sources

The research conducted by DT and BG on this project was funded by NIH grants U19 MH114821 and RF1MH128875. Support for DT, DS, and AMG was provided through NIH grant R01 NS121761. AMG and AS received funding from NIH grants R01NS086981 and R21NS121806.

## Notes

### Competing Interest Statement

The authors have declared no competing interest.

